# Sex-specific relationships between stress coping and avoidance behavior

**DOI:** 10.64898/2025.12.31.697182

**Authors:** Kailyn M. Price, Abigail M. Polter

## Abstract

While the experience of stress is ubiquitous, the risk of developing stress-linked conditions such as anxiety and depression is related to maladaptive stress responses. In order to probe the relationship between stress coping, sex, and stress-linked behavioral outcomes, we exposed male and female mice to subchronic variable stress (SCVS) and measured the correlation between coping during the tail suspension stressors (TSS) of SCVS and avoidance behavior in the EPM. We found that females engage in more active coping, and there were no sex differences in avoidance or locomotor behavior in the EPM after stress. However, we found that greater active coping predicted greater avoidance in females, but less avoidance in males. The results demonstrate that coping strategies are dynamic across time in males and females, but the relationships between avoidance and coping strategy dynamics are sex-biased.

**Plain English Summary:** The selection of stress coping strategies is an important component of the stress response that can impact behavior after stress. Stress coping strategies and behavior after stress can both be sex-biased, but the relationships between them are unclear. SCVS is a paradigm that is used to study sex differences in behavior and physiology because females are specifically vulnerable to SCVS. We recorded behavior during two stressors in the SCVS paradigm and found opposite relationships between coping behavior and avoidance behavior after stress in males and females, even though males and females exhibit similar dynamics in coping behavior and similar avoidance behavior after stress. These results demonstrate that sex is an important variable in the relationship between coping strategies during stress and behavior after stress.

## Introduction

All living organisms experience stress, broadly defined as challenges to physical or emotional homeostasis. Stress often carries a colloquially negative connotation, but it is ubiquitous and necessary for learning and adaptation [1, 2]. However, stress can precipitate maladaptive outcomes when it surpasses the adaptive capacities of an organism due to the chronicity, intensity, or perception of the stress. In humans, chronic stress exposure can increase the risk of developing multiple psychiatric disorders including depression, anxiety, and substance use-related conditions. Across organisms, stress initiates a cascade of physiological mechanisms and behavioral responses.

Among these responses are behavioral strategies known as stress coping, which permit the removal, mitigation, and adaptation to a stressor such that the stress response may be primed for more efficient responses in the future [3–5]. Stress coping strategies vary between conspecifics and across contexts, and some stress coping choices promote adaptation while others may be acutely or chronically maladaptive. The appraisal of stress and selection of coping strategy is influenced by a range of intrinsic factors including genetics, early life environment, and circulating hormones, which interact with real time stimuli and shared stress response mechanisms [2, 3]. Consequently, whether or not stress coping strategies are adaptive depends on a dynamic interplay of numerous factors including the intensity, frequency, and environmental context of the stressor, as well as the animal’s physical abilities and limitations [6–8]. For this reason, the investigation of stress coping strategies can aid the understanding of factors that drive individual differences in vulnerability and resilience for developing stress-linked conditions.

Epidemiological and preclinical data consistently demonstrate that sex and gender are key contributing factors in vulnerability for developing stress-linked conditions. The global prevalence of depression and anxiety-related conditions in women is two to three times greater than the prevalence in men, which likely results from a convergence of genetic, developmental, neurobiological, psychosocial, and cultural factors [9–12]. Latent cognitive processes, social support seeking, and anger-related traits are examples of gender-biased coping strategies that can directly contribute to the risk of experiencing new or recurrent depressive episodes [13, 14].

Sex-specific stress coping responses and vulnerability have also been observed in model organisms. Female rats exhibit greater corticosterone release and greater struggling behavior over multiple restraint stress sessions [15] and female mice are more susceptible to chronic mild stress as measured by greater immobility in the forced swim test and reduced population activity in the ventral tegmental area (VTA) after stress [16]. However, in conditioned fear contexts, females adopt the sex-biased strategy of darting, and darting females show reduced freezing during fear extinction [17]. These studies suggest that while females can engage in coping behavior during inescapable stressors that reflects reduced habituation to stress and impaired adaptation in stress pathways, they can also adopt specific behavioral strategies that promote adaptation. Thus, the role of behavioral strategies in adaptive responses is both sex and context-dependent.

The role of sex in stress-related neurobiological mechanisms and behavioral outcomes of stress has received increasing attention in recent years [18–22]. Paradigms that can model convergent and divergent sex-dependent mechanisms across behavioral and physiological endpoints are critical in advancing the understanding of links between stress, sex, physiology, and maladaptive behavioral outcomes. The subchronic variable stress (SCVS) paradigm models such divergence. This paradigm results in a range of sex-biased behavioral outcomes such as increased avoidance and anhedonia as well as physiological changes, including higher corticosterone release and changes in neuronal activity and gene expression across reward and limbic circuitry [23–30]. SCVS therefore reliably alters post-stress outcomes and physiology in a sex-dependent fashion, making it a robust platform for investigating whether sex-dependent coping strategies can lead to sexually divergent behavioral outcomes.

Tail suspension stress (TSS) is one of the three hour-long inescapable stressors employed during the SCVS paradigm. The tail suspension test (TST) was initially developed as a counterpart to the forced swim test (FST) that increased sensitivity for detecting anti-depressant effects of pharmacological treatments [31]. In its 6-minute form, greater immobility in the TST is typically seen as maladaptive-indicative of behavioral despair or overly passive responses. However, over a prolonged stressor, immobility is likely to be dynamic as animals respond to the repeated experience of unsuccessful escape attempts and balance the high energy cost of sustained struggling against the drive to escape. Given the female-specific vulnerability to SCVS, we hypothesized that females and males would display distinct patterns of coping during the TSS phases of stress, and that this behavior may predict post-stress behavioral avoidance. To test this hypothesis, we recorded stress coping behaviors during the TSS sessions of SCVS and measured their relationship with exploratory behavior in the EPM to examine relationships between sex, stress coping, and avoidance.

## Methods

### Animals

All experiments were conducted in accordance with National Institutes of Health Guidelines for the Care and Use of Laboratory Animals and approved by the Institutional Animal Care and Use Committee at The George Washington University. Male and female C57BL/6J mice were purchased from the Jackson Laboratory (#000664) or bred in-house. Mice were housed in groups of 3-5 in a temperature and humidity-controlled facility with *ad libitum* access to food and water on a 12:12 light/dark cycle for the duration of the experiment.

#### Subchronic variable stress

SCVS was performed as previously described [26, 29]. Briefly, 8 to 11-week-old male and female mice were exposed to one hour of foot shock, tail suspension, or restraint stress which alternated and repeated once over 6 days. On the first and fourth days, 100 0.5 mA foot shocks were randomly dispersed over one hour in a sound attenuated Coulbourn box. On the second and fifth days, mice were suspended by the tail with tape approximately 45 centimeters over the benchtop for 1 hour and behavior was video recorded at 30 fps. A lightweight tube was passed over the tail of the mouse to reduce tail climbing. On the third and sixth days, mice were placed in a ventilated 50 mL conical tube inside of their home cages for 1 hour. Males and females did not make physical contact with one another during stress or behavior.

#### TSS behavioral analysis

DBScorer, a MATLAB-based behavioral scoring software interface [32], was used to analyze struggling behavior in the TSS sessions. DBScorer reports immobility and struggling behaviors by calculating the change in the area occupied by the animal above a specified threshold between video frames. When the change in area does not exceed threshold, the animal is counted as immobile. Videos were analyzed in 10-minute bins across each tail suspension stressor, excluding the first 60 seconds of the first 10 minutes of stress. Each video was analyzed with blur image 0.1, 0.8% area threshold, 0s time threshold, and 60s time bin. Immobility behavior is reported in the text as the percent of time spent immobile between 0 and 100, where 0 is sustained struggling and 100 is full immobility. We also report immobility bouts, which is the number of times that the animal stopped struggling.

#### Elevated Plus Maze

Mice were acclimated to the testing room for 30 minutes before testing. Mice were placed in the center zone of a gray maze with their head facing the open arm opposite the experimenter and allowed to freely explore the maze for 6 minutes. The center zone was illuminated at 116 Lux. Opposing arms were 5.5 cm x 35 cm, raised 48 cm from the floor. Walls of closed arms were 15 cm high. Animals that fell off of the maze (n = 1 male, 1 female) were immediately placed back on the maze to complete testing but were excluded from analysis. Behavior was video recorded and analyzed with Any-maze software.

#### Statistical analysis

Data is reported as mean ± SEM. Statistical analyses were performed with GraphPad Prism 10.6.1. Outliers were not removed from the data set, the only excluded animals were those that fell off the EPM. For datasets that did not meet assumptions of normal distribution, non-parametric statistical tests were used. Statistical test and sample size details are indicated in each figure legend. Statistical significance level was set at p < 0.05 for all analyses.

## Results

### Females engage in more active coping during tail suspension stress

We first investigated whether males and females exhibited similar patterns of stress coping during the tail suspension sessions of SCVS. Average immobility scores were lower in females than males (main effect of sex, F_1, 56_ = 7.00, *p* = 0.011; Figure 1A), and higher during the second tail suspension session (main effect of session, F_1, 56_ = 32.78, *p* < 0.0001), but there was no significant sex x session interaction (F_1, 56_ = 3.76, *p* = 0.057). We also assessed the number of immobility bouts (Figure 1B), and found a significant main effect of sex (F_1, 56_ = 6.15, *p* = 0.016), but no significant effect of session (F_1, 56_ = 3.01, *p* = 0.088) or sex x session interaction (F_1, 56_ = 3.86, *p* = 0.055). This data suggests that during both TSS sessions, females are making more transitions between coping states, but are spending less time immobile between struggling bouts.

**Figure 1.**
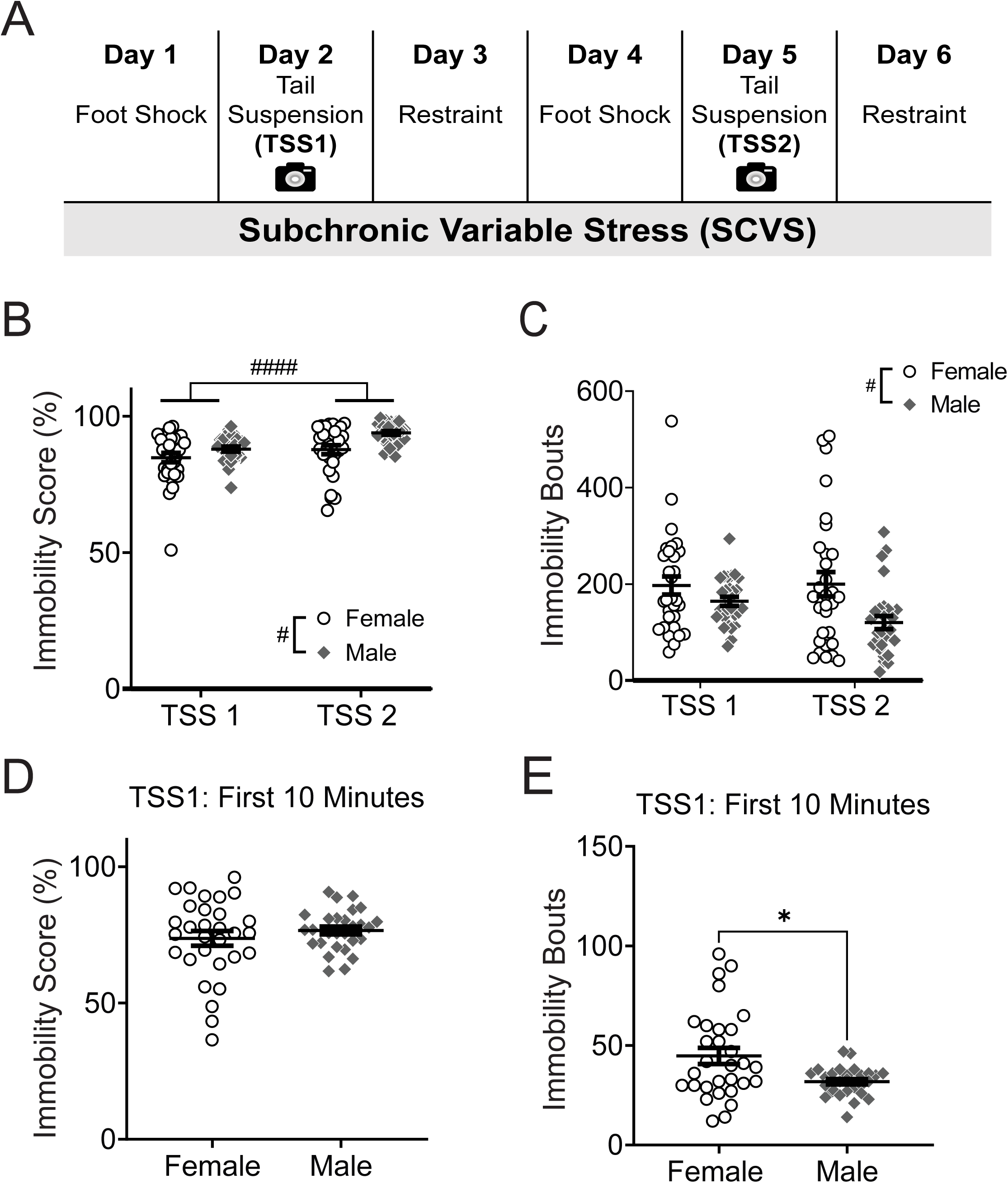
Stress coping behavior during the tail suspension phases of SCVS is sex-dependent. **A.** Experimental schematic for subchronic variable stress (SCVS). Behavior was video recorded during the first and second tail suspension sessions (TSS1 and TSS2). **B** Average immobility score and **C** total immobility bouts during TSS1 and TSS2. **D** Immobility score during the first 10 minutes of TSS1. **E** Total immobility bouts during the first 10 minutes of TSS1. # indicates significant main effect: # p < 0.05, ## p < 0.01, ### p < 0.001, #### p < 0.0001. Pairwise comparisons were performed with Mann-Whitney U test. N=28 males, 30 females. * p<0.05, ** p<0.01.

The tail suspension test, a classic test of stress coping strategy and antidepressant responses, is traditionally 6-10 minutes in duration [31, 33]. In order to test whether we would have detected sex differences in coping behavior in this duration of test, we compared immobility scores and bouts during the first 10 minutes of the first TSS session. We found no significant sex difference in immobility score (*U* = 392.5, *p* = 0.67; Figure 1C), but did find that there were more immobility bouts in females (*U* = 279, *p* = 0.028; Figure 1D). These results further suggest that sex differences emerge in immobility behavior over the one-hour tail suspension sessions and would not be detectable by measuring the immobility score in the first 10 minutes of stress alone.

While average immobility scores and total immobility bouts illustrate activity over the entire stressor, they do not demonstrate behavioral changes during prolonged stress, an important measure of learning and adaptation. For this reason, we examined the change in immobility scores and immobility bouts within 10-minute bins over the duration of both tail suspension sessions (Figure 2A-D). There was a significant main effect of time on the immobility score during the first TSS session (F_2.94, 164.8_ = 46.34, *p* < 0.0001; Figure 2A) but no main effect of sex (F_1, 56_ = 2.46, *p* = 0.12) or sex x time interaction (F_2.94, 164.8_ = 0.14, *p* = 0.94). Immobility scores increased during each 10-minute bin after the first 10 minutes (Figure 2A). There was also a significant main effect of time on immobility bouts during the first TSS session (F_3.45, 193.0_ = 16.36, *p* < 0.0001; Figure 2B), but no main effect of sex (F_1, 56_ = 2.33, *p* = 0.13) or sex x time interaction (F_3.45, 193.0_ = 2.01, *p* = 0.10). The number of immobility bouts decreased over time, suggesting that animals made fewer transitions between coping states and spent most of their time immobile. During the second TSS session, there was a main effect of sex on immobility score (F_1, 56_ = 11.13, *p* = 0.0015; Figure 2C), but not time (F_3.85, 215.5_ = 1.56, *p* = 0.19) or sex x time interaction (F_3.85, 215.5_ = 0.27, *p =* 0.89), with females spending less time immobile than males for the duration of the stressor. However, there was a significant main effect of both sex (F_1, 56_ = 7.46, *p* = 0.0084; Figure 2D) and time (F_2.69, 150.7_ = 4.66, *p* = 0.0053) on the number of immobility bouts in TSS2, but no significant sex x time interaction (F_2.69, 150.7_ = 0.072, *p* = 0.97). Further analysis demonstrated that the session day significantly contributed to the magnitude of immobility score change (F_1, 56_ = 74.78, *p* < 0.0001; Figure 2E) and immobility bout change (F_1, 56_ = 40.51, *p* < 0.0001; Figure 2F) within animals. However, changes in immobility score and immobility bouts over time did not differ by sex (immobility score: F_1, 56_ = 0.25, *p* = 0.62, immobility bouts: F_1, 56_ = 1.09, *p* = 0.30) or a sex x session interaction (immobility score: F_1, 56_ = 0.0038, *p* = 0.95, immobility bouts: F_1, 56_ = 1.92, *p* = 0.17). These results suggest that changes over time in immobility behavior occur predominantly over the first exposure to TSS and are similar in males and females, therefore the sex differences in average immobility scores and bouts in the second TSS session result from sustained coping strategies selected at the beginning of stress, not the rate of adaptation to stress.

**Figure 2.**
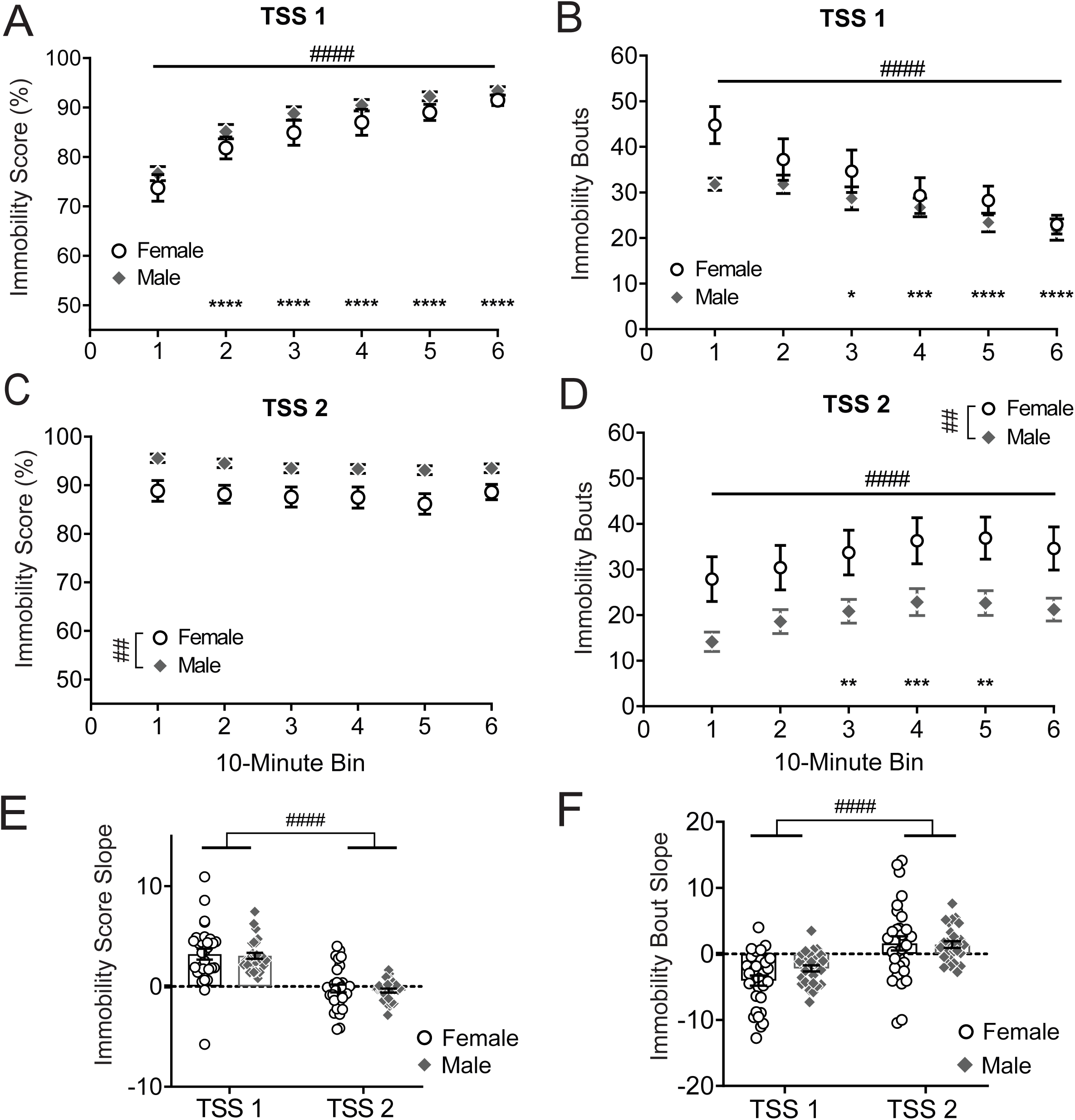
Stress coping behavior changes within and between tail suspension sessions. **A.** Immobility score and **B** immobility bouts across TSS1, within 10-minute bins. **C.** Immobility score and **D** Immobility bouts across TSS2, within 10-minute bins. Two-way ANOVA followed by Dunnett’s multiple comparisons test where significant main effects of time were present. **E** Slopes of immobility score and **F** slopes of immobility bouts over time for each animal. Two-way ANOVA. # indicates significant main effect: # p < 0.05, ## p < 0.01, ### p < 0.001, #### p < 0.0001. * indicates significant pairwise difference in comparison to first 10 minutes. * p < 0.05, ** p < 0.01, *** p < 0.001, ****p < 0.0001. N = 28 males, 30 females.

### Coping behavior during stress predicts avoidance

Given the known roles of sex in risk appraisal, risk taking, and emotional reactivity [34, 35], we sought to determine whether the sex-dependent coping behaviors discovered in the TSS sessions would be associated with post-stress behavior in the EPM. We first assessed whether there were sex differences in exploratory behaviors in the EPM. We found no sex differences in total distance traveled (*t*_38_ = 0.79, *p* = 0.43; Figure 3A), open arm entries (*t*_38_ = 0.50, *p* = 0.62; Figure 3B), open arm time (*t*_38_ = 1.45, *p* = 0.15; Figure 3C), or open arm ratio (*t*_38_ = 1.63, *p* = 0.11; Figure 3D).

**Figure 3.**
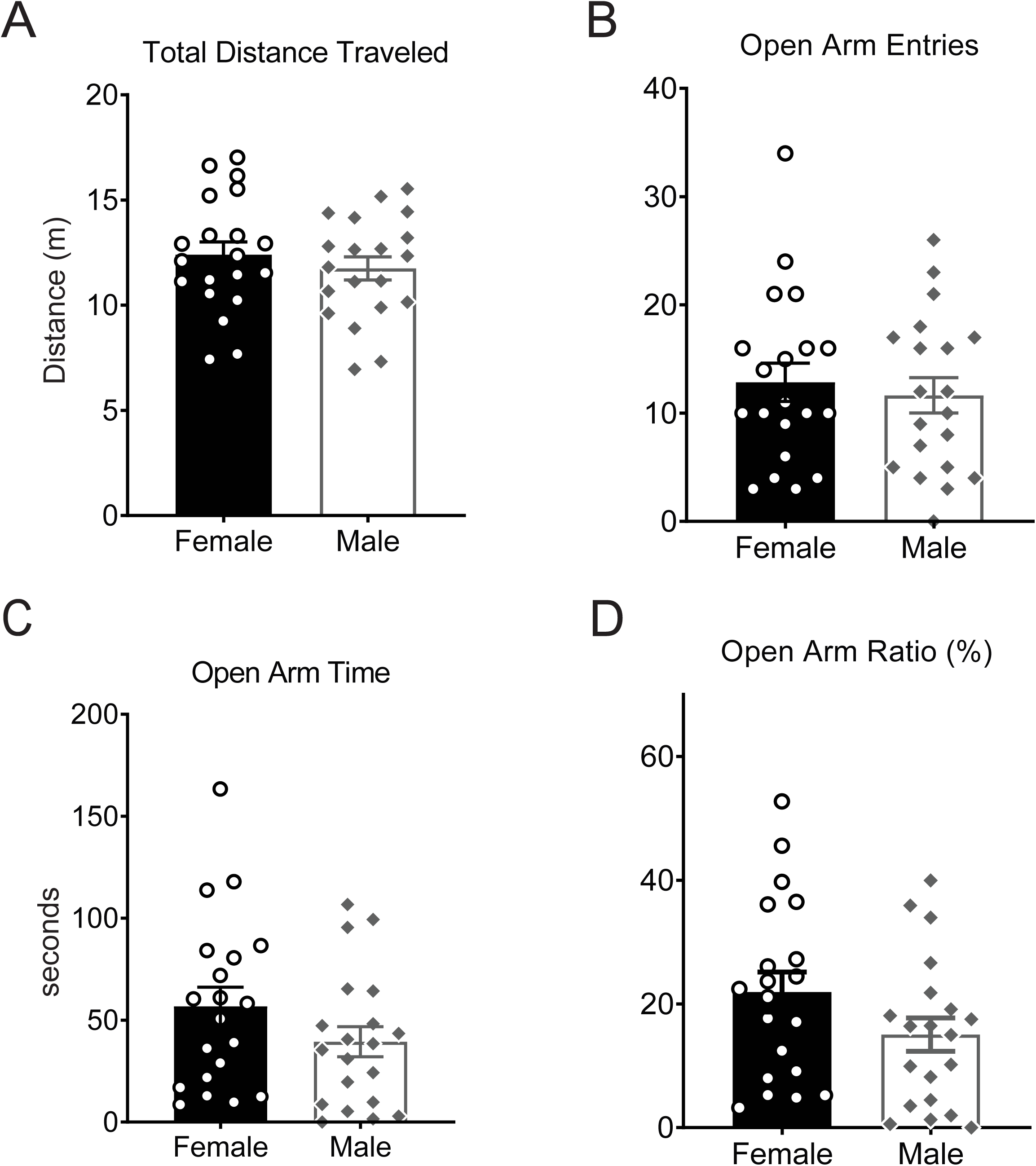
No sex differences in avoidance behaviors in the elevated plus maze. After SCVS, **A**. total distance traveled (males: 12.40 ± 0.61 m, females: 11.75 ± 0.55 m) in the elevated plus maze. **B**. open arm entries (males: 11.65 ± 1.64, females: 12.85 ± 1.77, **C**. open arm time (males: 39.40 ± 7.39 s, females: 56.78 ± 9.40 s) and **D.** open arm ratio (males: 15.05 ± 2.70%, females: 21.92 ± 3.24%) were measured in males and females. Open arm ratio = [(open arm time/ open arm time + closed arm time) * 100]. N = 20 females, 20 males.

One possible contributor to variability within males and females in avoidance behaviors after stress may be sex differences in the appraisal of and responses to stress that directly contribute to the appraisal of threatening contexts after stress. In order to test relationships between coping behavior and avoidance, we performed simple linear regressions on average immobility scores and open arm ratios. In males, higher immobility scores during the first TSS session predicted less open arm time, or greater avoidance (Figure 4A; R^2^ = 0.29, *p* = 0.014). In the second TSS session, males’ immobility scores and open arm ratios showed a similar relationship but did not reach significance (Figure 4C; R^2^ = 0.14, *p* = 0.10). In females, however, greater immobility scores predicted a higher OA ratio, or *less* avoidance in both the first (Figure 4B; R^2^ = 0.27, *p* = 0.02) and second (Figure 4D; R^2^ = 0.43, *p* = 0.0017) TSS sessions. The slopes of the immobility score and OA ratio regression were significantly different between males and females for the first (F_1,36_ = 12.11, *p =* 0.0013) and second (F_1,36_ = 7.90, *p =* 0.008) tail suspension sessions. To control for locomotor activity, we performed simple linear regression analyses of distance traveled in the EPM and immobility scores during both tail suspension sessions (Figure 4E-F), and found no significant relationships in males (TSS1: R^2^ = 0.062, p = 0.29, TSS2: R^2^ = 0.13, *p* = 0.12) or females (TSS1: R^2^ = 0.00081, p = 0.91, TSS2: R^2^ = 0.0047, *p* = 0.77). Together, these results demonstrate that the relationships between behavior during stress and avoidance behavior are both sex and time dependent.

**Figure 4.**
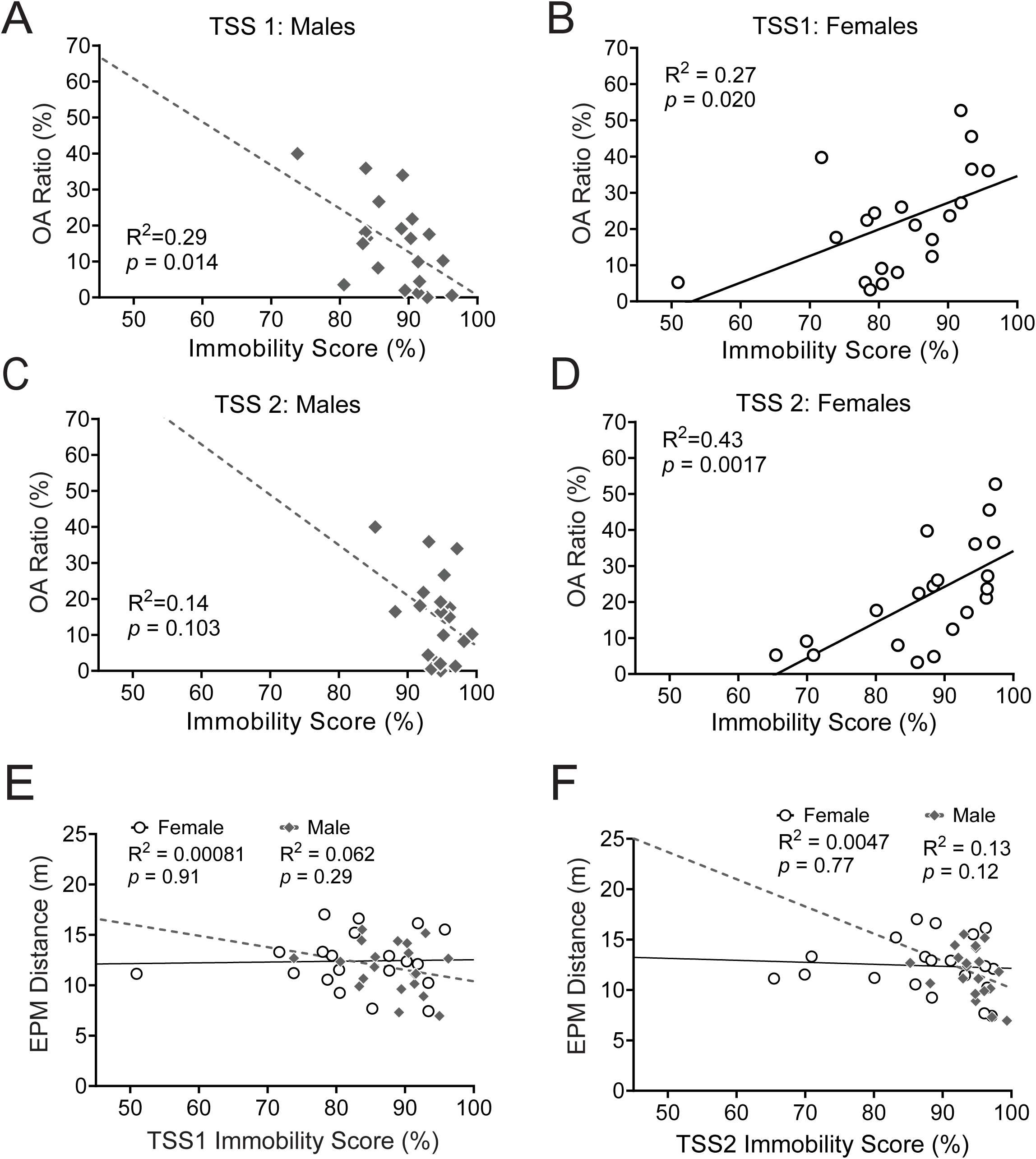
Relationships between stress coping and post-stress avoidance behavior are sex-specific. Simple linear regressions between OA ratio and immobility score during TSS1 in males **A** and females **B**, and OA ratio and immobility score during TSS2 in males **C** and females **D**. Simple linear regression of distance traveled in the elevated plus maze and immobility score during TSS1 **E** and TSS2 **F**. R^2^ and p-values listed on graph inset. N= 20 females, 20 males.

### Relationships between stress coping and avoidance are sex-dependent

Given the relationships between avoidance and behavior in the TSS, we tested the collinearity of selected behavioral measures across the tail suspension stressors and avoidance behavior (Figure 5). We found that in females, immobility score during the first 10 minutes of TSS1 was positively correlated with the average TSS1 immobility score (r = 0.60, *p* = 0.005), TSS2 immobility score (r = 0.78, *p* < 0.0001), and OA ratio (r = 0.61, *p* = 0.005), demonstrating that coping behavior in the first 10 minutes of TSS could predict behavior across both tail suspension sessions in addition to post-stress avoidance behavior. Immobility scores during the first 10 minutes of the first TSS stressor were negatively correlated with the slope of the score over the session (r = − 0.91, *p* < 0.0001), indicating that lower immobility at the start of the session was correlated with a greater change in coping over the first stress session. Interestingly, the immobility score slope in females was inversely correlated with OA ratio (r = −0.59, *p* = 0.006), which suggests that a higher rate of coping style change in females predicts more avoidance after stress. This is likely explained mostly by the inverse relationship between immobility in the first 10 minutes of stress and immobility slope, as females who start at a lower immobility score have a higher change in their coping score over time, and the immobility score in the first 10 minutes alone predicts avoidance after stress.

**Figure 5.**
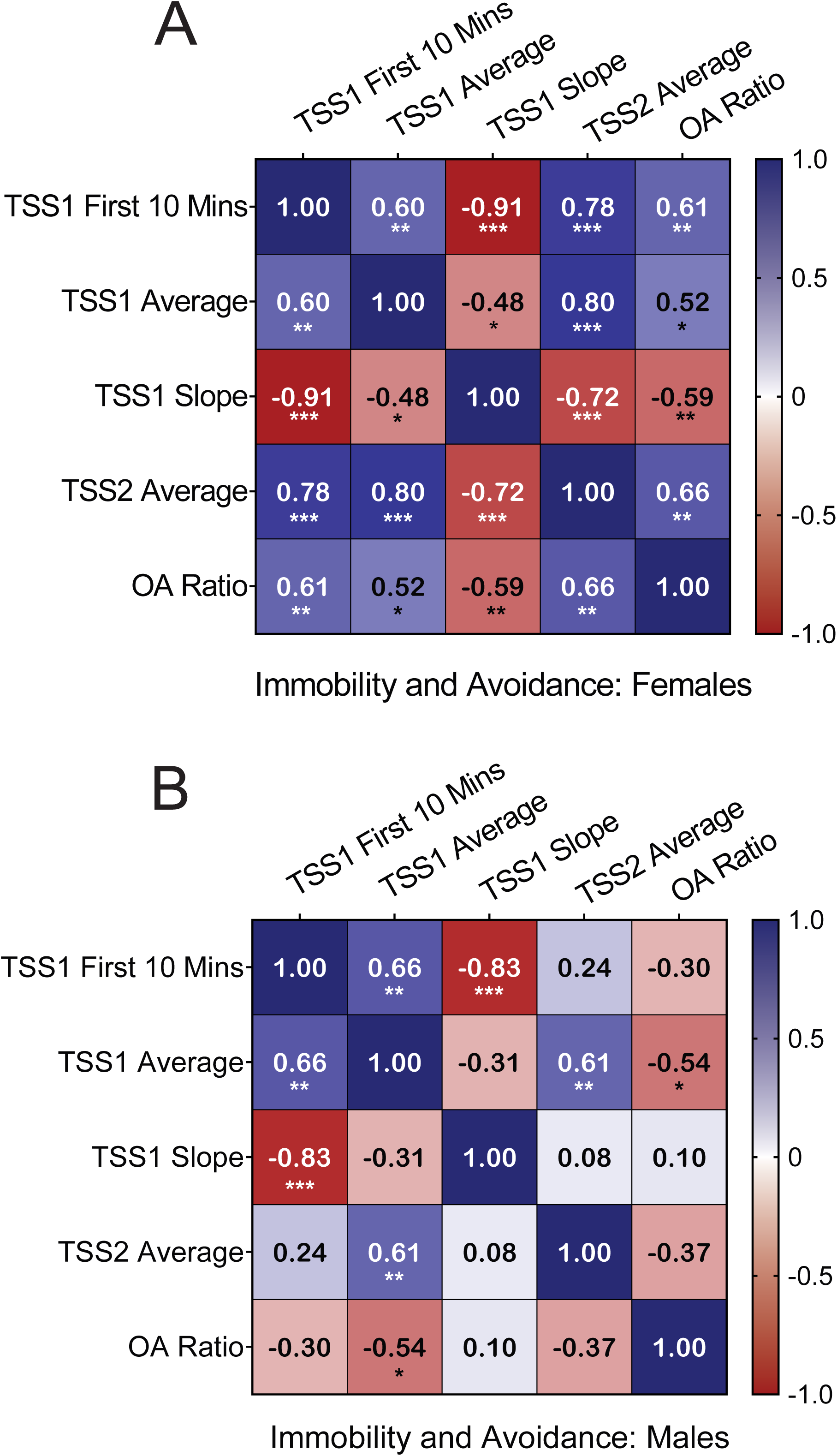
Covariance between immobility measures and avoidance behaviors is sex-dependent. Pearson r correlation coefficients between the first 10 minutes of TSS1, TSS1 average immobility score, TSS1 immobility slope, TSS2 average immobility score, and open arm ratio in females **A** and males **B**. N= 20 females, 20 males. * p < 0.05, ** p < 0.01, *** p < 0.001, ****p < 0.0001.

TSS1 and TSS2 immobility scores were positively correlated in males (r = 0.61, *p*= 0.005) and females (r = 0.80, *p* < 0.0001), which suggests that coping choices within animals are consistent between stress sessions regardless of sex. However, in males, while immobility in the first 10 minutes of TSS1 was positively correlated with the average immobility score during TSS1 (r = 0.66, *p* = 0.002) and inversely correlated with the slope of TSS1 immobility score (r = −0.83, *p* < 0.0001), it did not significantly correlate with coping behavior in TSS2 (r = 0.24, p = 0.31) or with OA ratio (r = −0.30, *p* = 0.20) unlike the observation in females. Simple linear regression revealed significant sex differences between the immobility score in the first 10 minutes of TSS1 and OA ratio (F_1, 36_ = 7.68, *p* = 0.0088), as well as the TSS1 immobility score slope and OA ratio (F_1, 36_ = 4.53, *p* = 0.040). Taken together, these results suggest that the relationships between behavioral choices at the beginning of stress and behavior across both stress sessions, and the extent to which those behavioral choices predict post-stress avoidance, are sex-dependent. For females, behavior at the beginning of stress predicts the behavioral profile across both stress sessions and is sufficient to predict avoidance after stress. In males, however, behavior at the beginning of stress is only predictive of the behavioral profile during the first stress session. This suggests that while males and females engage in similar magnitudes of behavioral flexibility as measured by the change in immobility over TSS, females select a set of behavioral choices at the beginning of stress that are sustained across multiple stress sessions and predict avoidance after stress, while the initial behavioral strategies in males are not necessarily sustained across both sessions and do not predict avoidance.

We considered one possible source of behavioral variability contributing to sex differences in coping behavior-adoption of tail climbing. We assessed whether animals tail climbed at any point during TSS sessions and found that 46.67% of females engaged in tail climbing at some point during the first tail suspension session and 36.67% engaged in tail climbing during the second tail suspension session, while only 7.14% of males engaged in tail climbing during either TSS session (Figure 6). The proportion of tail climbing was significantly different between males and females during both the first (*p* = 0.0010) and second (*p* = 0.011) TSS sessions (Fisher’s exact test; Figure 6A-D). Within females, there was a significant main effect of tail climbing status on immobility score, with tail climbing females having a significantly lower immobility score than non-tail-climbers (F_1, 56_ = 64.43, *p* < 0.0001; Figure 6E). Because DBScorer does not differentiate between head down and tail climb struggling bouts, it is unclear what proportion of each total immobility score is attributable to tail climbing vs. head down struggling. However, it is clear that in this configuration tail climbing does not result in a broadly immobile phenotype during tail suspension and that it is a sex-biased strategy that may contribute to adaptation during stress.

**Figure 6.**
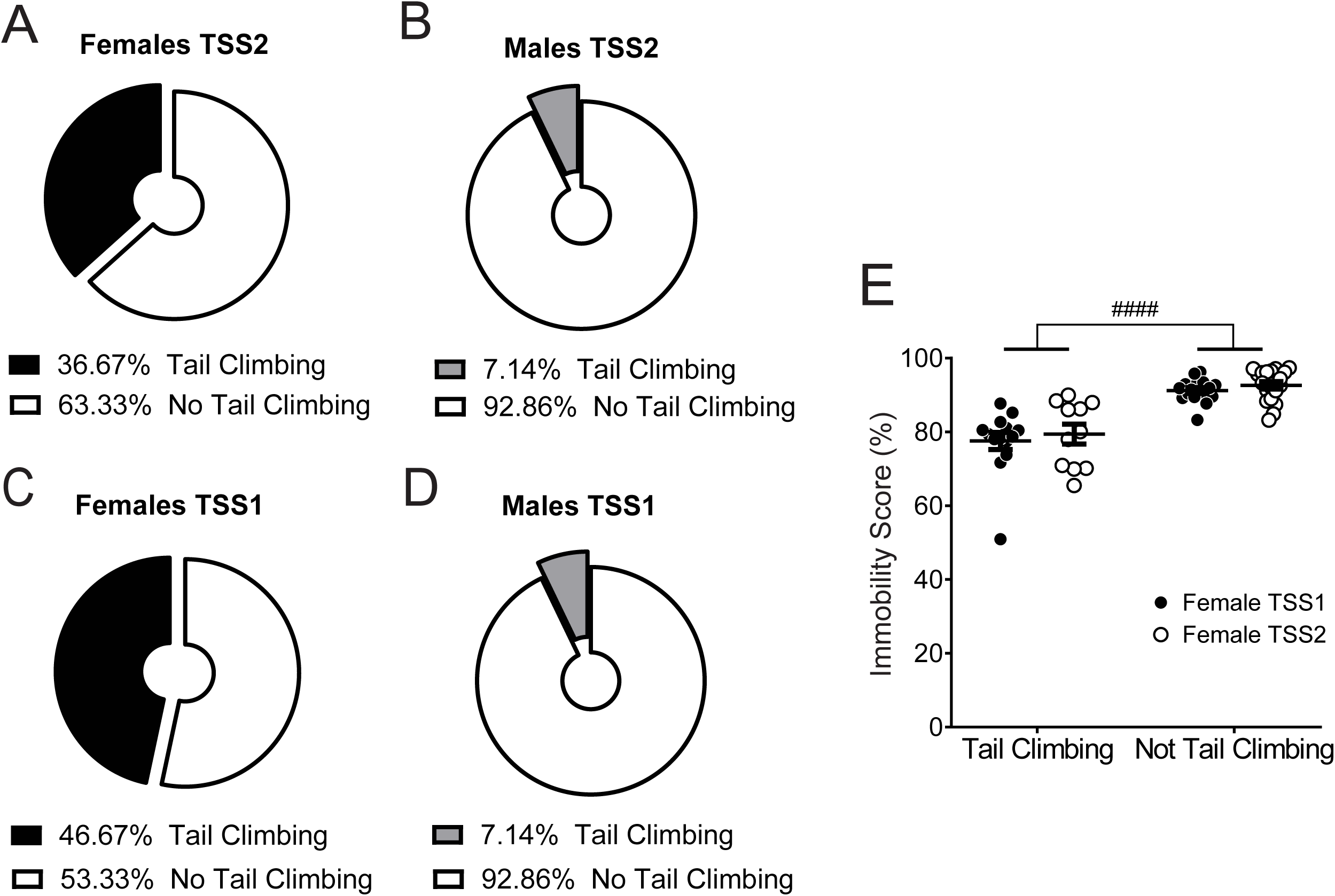
Tail climbing rates are greater in females during tail suspension stress. Proportions of tail climbing at any point during TSS1 in females **A** and males **B** and during TSS2 in females **C** and males **D**. N = 28 males, 30 females. **E**. Immobility scores are higher in females who do not tail climb at any point during TSS. Two-way ANOVA. #### indicates main effect, p < 0.0001. N = 11-19/group.

## Discussion

In this study, we tested whether sex differences in coping strategies emerge during a repeated stressor, and if they are associated with avoidance behaviors after stress. Active coping is often seen as a beneficial or adaptive choice that promotes resilience after stress, whereas passive coping indicates despair or a “depressive-like” phenotype [36]. However, in an inescapable stressor, the choice to sustain a coping strategy that expends considerable energy may reflect a failure to learn. Sustained active coping may lead to pathophysiological plasticity in neural circuitry that contributes to avoidance, reward, motivation, and aversion. This is particularly important in the SCVS paradigm where animals are exposed to a series of inescapable stressors and must repeatedly select strategies that promote adaptation across each stressor. We focused on behavior in the TSS sessions of SCVS, but it is likely that behavior is influenced by previous and ongoing experience across each stressor and would differ from behavior during isolated tail suspension tests. These prior and ongoing experiences may be important for the observed sex differences in the relationship between coping and avoidance after stress, specifically given observed sex differences in behavioral strategies during other inescapable stressors [15, 17].

One strength of using the SCVS paradigm to ask this question is that each stressor is repeated, which allows comparison of behavior during repeated stress sessions to ascertain changes in coping strategy. Our data demonstrates that sex differences in coping strategy are a function of both time and repeated exposure to stress. While the choice of coping strategy changed over the duration of the first stress session in both males and females, sex differences were observed in overall immobility score only in the second tail suspension session, where the choice of strategy at the beginning of the stressor was sustained over the duration of the session. This suggests that there may be sex differences in mechanisms that support sustained motivation and learning across repeated inescapable stress sessions.

Our study demonstrates that relationships between avoidance behavior and coping behavior across stress sessions are a meaningful sex-dependent outcome of SCVS and predict behavioral variability within each sex. Despite no sex differences in the overall immobility score during the first TSS session or EPM open arm ratio, higher immobility scores predicted greater avoidance in males but lower avoidance in females. On the second TSS session, when females engaged in more active coping than males, higher immobility scores were only a significant predictor of avoidance in females, but not males. Importantly, some studies have identified sex differences in baseline locomotion of mice and rats during the EPM, which can confound interpretations of exploratory behavior [37, 38]. However, our study shows that there is no relationship between the immobility score and locomotion in the EPM. This suggests that the relationship between coping strategy during tail suspension and avoidance is not reducible to sex-biased trends in activity levels.

Assays like the tail suspension and forced swim tests often report total time spent immobile, or inversely, time spent on escape-oriented behaviors. This data suggests, however, that the number of transitions between immobility and escape-oriented behavior may reveal important information that total time in each state may not capture. During the first 10 minutes of the first TSS session, there was no sex difference in the overall immobility score, but females made significantly more transitions between immobility and struggling as measured by the number of immobility bouts. The differences between immobility scores and immobility bouts may reflect a distinction between action initiation for short escape-oriented bouts in comparison to the motivational vigor required for sustaining longer struggling bouts [39, 40].

Researchers utilizing the tail suspension test for screening antidepressants and measuring stress-induced behavioral changes often highlight the challenge of high tail climbing rates in C57BL/6J mice, and many exclude animals who tail climb [33, 41–43]. We did not apply this exclusion criterion in our study for several key reasons. Tail climbing is a coping strategy-it does not permit the animal to escape and requires energy expenditure in a similar way to head-down escape-oriented motion. Importantly for our study, we also found that tail climbing behavior is sex-biased. Nearly half of females but almost no males exhibit tail climbing at some point during the tail suspension stressor. Removing tail climbing animals would introduce a sex-biased exclusion criterion that would preclude a full assessment of how sex-biased coping strategies contribute to behavioral outcomes. It is possible that the vestibular and proprioceptive experience in an entirely head-down position is distinct and that repeated tail climbing promotes greater motivation to struggle, both of which may be important for understanding neural mechanisms engaged by struggling behavior. As Shansky and Murphy (2021) have emphasized, including females in studies warrants consideration that common behavioral endpoints based on assays that were originally tested exclusively in males may not reflect the range of meaningful behavioral strategies in females [21].

While the EPM is generally understood to test a conserved conflict between an aversion to open or elevated space and exploration, others argue that it is also testing thigmotaxis mediated by the somatosensory system, primarily in the closed arms of the maze [44]. Studies have identified sex differences in thigmotaxis across anxiogenic environments, which may contribute to sex differences in exploratory behavior in the EPM [45, 46]. Neural circuits that assess threat and support coping strategy selection during stress may be altered by repeated unsuccessful escape attempts during stress, which could directly inform future threat assessments in the EPM. These circuits are reliant upon sensory input and interoceptive signals during stress and motivated behavior, and in anxiogenic environments [47–51]. Thus, appraisal of risk and safety signals may be altered by coping choices to promote adaptation after chronic stress. Furthermore, each stressor was performed in groups with cage mates, so while animals could not make physical contact in TSS, it is likely that they can see, smell, and hear their cage mates during stress. Prior work has demonstrated that a physical barrier or being tested alone does not alter behavior in males during the 6-minute tail suspension test [52], but it is unclear whether this would be true in females, and whether it would apply in a prolonged stress session. Given the sex-specific roles of social stress on physiological and behavioral outcomes [18], future studies to test the role of social context and sensory cues during stress on sex-specific coping strategies and avoidance behaviors would be beneficial.

Neither coping strategies nor avoidance behavior can be interpreted under binaries-the more likely case is that an accumulation of maladaptive choices, particularly when those behaviors are sustained over time, can contribute to reduced fitness or pathophysiological states. Why the relationship between coping choices and avoidance would be sex-biased is likely attributable to differences in the evolutionary roles of escape behaviors during inescapable stress and approach behaviors, which engage overlapping neural circuitry [53]. Fluctuations in sex hormones across the estrous cycle and between animals of different social rank may contribute to baseline stress and anxiety levels, and act directly on limbic and striatal circuitry that drive threat appraisal and memory [54–57]. The BNST is a known substrate for both stress coping and avoidance behaviors [48, 49] with established sex differences in contributions to threat processing [58]. The locus coeruleus and ventral tegmental area, hubs for robustly stress-sensitive noradrenergic and dopaminergic circuitry respectively, exhibit structural and physiological sex differences that may also play direct roles in stress processing that are particularly relevant in convergent inputs to limbic and cortical regions [59]. Future studies exploring specific contributions of sex hormones and sex biased regions to learning and adaptation during stress coping could advance concepts of the roles of sex in stress coping and avoidance behavior.

## Summary/ Conclusions

Our study demonstrates that coping strategies are sex-specific and dynamic across a single stress session and between repeated stress exposures. Furthermore, coping strategies and the change in coping strategy predicts avoidance behavior in a sex-dependent fashion. While coping strategies are often not recorded and scored during stress, they may provide a key readout for the progression of behavioral changes across chronic stress paradigms, and may contribute to elucidating the engagement of mechanisms that promote divergent and convergent sex-specific mechanisms.

## Declarations

### Ethics approval and consent to participate

Not applicable

## Consent for publication

Not applicable

## Availability of data and materials

The datasets used and analyzed during the current study are available from the corresponding author on reasonable request.

## Author’s contributions

KP and AMP designed the study; KP collected and analyzed data; KP and AMP interpreted data; KP drafted the paper; AMP edited the paper.

## Acknowledgments

We would like to acknowledge Dr. Paul Marvar for use of the Coulbourn boxes.

## Author Information

Department of Pharmacology and Physiology, George Washington University School of Medicine and Health Sciences, Washington, DC 20037

